# Asymmetric gene flow across a desert contact zone in a riparian songbird

**DOI:** 10.64898/2026.02.04.703846

**Authors:** Ethan F. Gyllenhaal, Andrew B. Johnson, Lukas B. Klicka, Selina M. Bauernfeind, Matthew J. Baumann, Matthew L. Brady, Kevin J. Burns, Christopher C. Witt, Michael J. Andersen

**Affiliations:** Department of Biology, University of New Mexico, Albuquerque, New Mexico, USA; Museum of Southwestern Biology, University of New Mexico, Albuquerque, New Mexico, USA; Department of Biological Sciences, Texas Tech University, Lubbock, Texas, USA; Cornell University, Department of Computational Biology, Ithaca, New York, USA; Department of Biology, San Diego State University, San Diego, California, USA; School of Arts & Sciences, Peru State College, Peru, Nebraska, USA; Department of Biological Sciences and Biodiversity Collections, The University of Texas El Paso, El Paso, Texas, USA

**Keywords:** gene flow, phylogeography, genomics, Pleistocene refugia, hybrid population

## Abstract

Secondary contact is a key point in the speciation process, and fine-scale geography can shape its outcomes. This is especially true for species restricted to fragmented habitats, such as riparian corridors through arid regions. Here we examine the role of disjunct riparian habitat in shaping secondary contact in Bell’s Vireo (*Vireo bellii*), a North American songbird species that contains distinct eastern and western forms. We recovered a unique, discontinuous contact zone along the Rio Grande in New Mexico, where two populations with greater nuclear and mitochondrial genetic affinity for the eastern lineage are separated by a population with an affinity for the western lineage. This point of primarily western ancestry on the Rio Grande corresponded with a stretch where several intermittently flowing tributaries join from the west and may have acted as gene flow corridors. Using a combination of empirical analyses of divergence and diversity across the genome and population genetic simulations, we uncovered evidence of neutral, genome-wide admixture driving the genomic architecture of divergence, rather than evidence for local adaptation or selective sweeps. In sum, this genomic study showed us how fine-scale dispersal corridors can cause idiosyncratic patterns of admixture when habitat is limiting in zones of secondary contact.

## Introduction

Cases of secondary contact where divergent lineages meet and can exchange genes are key points in the speciation process and are strongly impacted by the geography in the region of contact (Hewitt 2001). Contact zones remain a classic model system for understanding introgression dynamics in divergent gene pools at the point of secondary contact (Mayr 1942, Shogren et al. 2024). Additionally, dynamics in these contact zones contribute fundamentally to our comprehension of speciation and are essential for predicting how populations will respond to anthropogenic change (Bontrager et al. 2019, Anderson et al. 2025). However, to do so we must have a strong understanding of how climate and geography shape the outcomes of secondary contact, particularly symmetry of introgression.

Most research on hybridizing taxa has focused on hybrid zones within continuous habitats (Barton 1983, Barton & Hewitt 1985, Wait & Peñalba 2025). However, geographic factors and spatially variable selection can lead to alternate patterns, such as mosaic hybrid zones (Harrison & Larson 2014, Curry 2015) and hybrid populations (Barrera-Guzmán et al. 2017, Andersen et al. 2021, Gyllenhaal et al. 2025a). In Nearctic birds, rapid post-glacial expansions have tended to produce continuous, clinal hybrid zones that are appropriate for cline-based modeling (Rising 1983, Swenson & Howard 2005, Walsh et al. 2020). This is best exemplified by suture zones where multiple hybrid zones overlap and allow comparative study and have significantly advanced our understanding of the genomic architecture and fitness consequences of secondary contact (Irwin et al. 2018, Mikkelsen & Irwin 2021, Rowher et al. 2023, Walsh et al. 2023, Wait & Peñalba 2025, DeRaad et al. 2025). However, traditional geographic cline methods are not easily applied to discontinuous hybrid zones, necessitating alternative approaches and models.

Habitat fragmentation—both natural and anthropogenic—can be one driver of discontinuous contact zones. Island biogeography has long informed studies of fragmented continental populations, particularly with respect to the species-area relationship (MacArthur & Wilson 1967, Lomolino & Perault 2001, Whittaker et al. 2005). Although historically emphasizing community composition, island biogeographic theory can also predict the asymmetry of dispersal—and thus gene flow—in island systems (Gyllenhaal et al. 2020). As with the species-area relationship, these models can also be applied to fragmented, island-like habitats on continents. Key predictions include larger fragments acting as stronger sources of migrants and fragments with a large "target size"—the width relative to the source of migrants—as likely recipients of migrants.

Asymmetric migration can range in impact from genetic swamping (Roberts et al. 2010, Rutherford et al. 2019) to beneficial adaptive introgression or maintenance of genetic health (Consuegra et al. 2005, Paz-Vinas et al. 2013, Tomasini & Peischl 2020, Pfennig 2021). While models of speciation commonly predict highly co-evolved gene networks, universally beneficial alleles may arise independently through mutation-order processes (Schluter 2009, Irwin et al. 2018). Past work has suggested that selection on universally beneficial alleles, sometimes facilitated by introgression post-divergence, best explains genomic divergence peaks (e.g., elevated F_ST;_ Cruickshank & Hahn 2014, Bay & Ruegg 2017, Irwin et al. 2018). However, these studies typically assume symmetric gene flow and assess divergence primarily through mean within-population diversity. In asymmetric adaptive introgression, we might expect to observe lower within-population diversity primarily in the donor population during allele sweeps into recipient populations (Bay & Ruegg 2017). Alternatively, asymmetric introgression of genomic regions with inherently reduced diversity due to low recombination rates could produce this skewed diversity pattern without invoking selective sweeps.

Riparian habitats across the arid Southwestern United States face threats from widespread habitat degradation, water diversion, droughts, and invasive species (Knopf et al. 1988, Sogge et al. 2008, Poff et al. 2011), making species inhabiting these ecosystems a research and conservation priority (Pilger et al. 2017, Turner et al. 2023). Southwestern New Mexico, particularly along the continental divide, corresponds roughly to the Cochise Filter Barrier, a well-known biogeographic barrier in desert taxa (Provost et al. 2018, Provost et al. 2021). Riparian habitats in this region, isolated by surrounding desert grasslands and scrub, may function similarly to islands, allowing the application of island biogeographic theory to model genetic exchange (Gyllenhaal et al. 2020, Gyllenhaal et al. 2025b). This can also mean that relatively large but isolated rivers like the Rio Grande could act as net sinks to gene flow from adjacent drainage systems with less water more overall riparian habitat.

Bell’s Vireo (*Vireo bellii*) is an ideal species for examining these processes. This songbird inhabits shrubby environments throughout the central and southwestern U.S. and northern Mexico. In the southwestern United States, it is confined primarily to riparian zones. In New Mexico, it is notably restricted to larger river drainages with enough water to maintain dense riparian habitat (Kus et al. 2022). Populations across its range have declined significantly in recent decades, resulting in its classification as ‘threatened’ in New Mexico and several other states (Sauer et al. 2015). Four modestly differentiated subspecies span its longitudinal distribution, from east to west: *V. b. bellii*, *V. b. medius*, *V. b. arizonae*, and *V. b. pusillus* (Kus et al. 2022). The two eastern subspecies tend to be characterized by a brighter hue to their flank coloration, while the western two tend to be duller. *Vireo b. medius* occurs in much of Texas and is thought to be the subspecies in eastern and central New Mexico within the Pecos River and Rio Grande drainages (Hubbard 1970). *Vireo b. arizonae* is common across southern Arizona and is the subspecies thought to occur in southwestern New Mexico in the Gila River drainage (Hubbard 1971).

Previous genetic studies found substantial mitochondrial and nuclear divergence between eastern and western subspecies pairs, suggesting species-level differentiation (Klicka et al. 2016). These analyses, complemented by species distribution modeling, inferred historical refugia in portions of the Rio Grande in what is now West Texas, facilitating rapid post-glacial expansion along associated riparian corridors. Given this historical context and deep genetic divergence, Bell’s Vireo presents an excellent system to explore how fragmented riparian habitats shape population structure and gene flow in the Desert Southwest. We hypothesize that that the uniquely fragmented habitat in this region resulted in a discontinuous hybrid zone, characterized by complex admixture patterns along the Rio Grande Valley. Leveraging a whole-genome dataset, we test this hypothesis and examine the heterogeneity in genomic divergence within this system.

## Methods

### Sampling

We sampled 44 individuals of *Vireo bellii* using muscle tissue (Table S1). Of these, 20 were from New Mexico, 10 from Arizona (*V. b. arizonae*), 13 from Texas (putative *V. b. medius*; 11 from West Texas and 2 from the Lower Rio Grande), and 1 from Missouri (*V. b. bellii*). All Arizona and Missouri samples, as well as 10 Texas samples were previously the subjects of a RADseq study (Klicka et al. 2016). Within New Mexico, samples included 11 from the Rio Grande Valley (two sampling sites, three from Sevilleta National Wildlife Refuge and eight from north of Elephant Butte Lake), and three each from the Mimbres, Pecos, and Gila rivers (Figure 1). All samples originated from vouchered museum specimens archived in five institutions (Table S1). Sampling maps were generated with the sf v1.0-16 (Pebesma 2018) package implemented in R v 4.3.0 (R Core Team 2023), leveraging shapefiles from HydroBASINS and HydroRIVERS (Lehner & Grill 2013).

**Figure 1:**
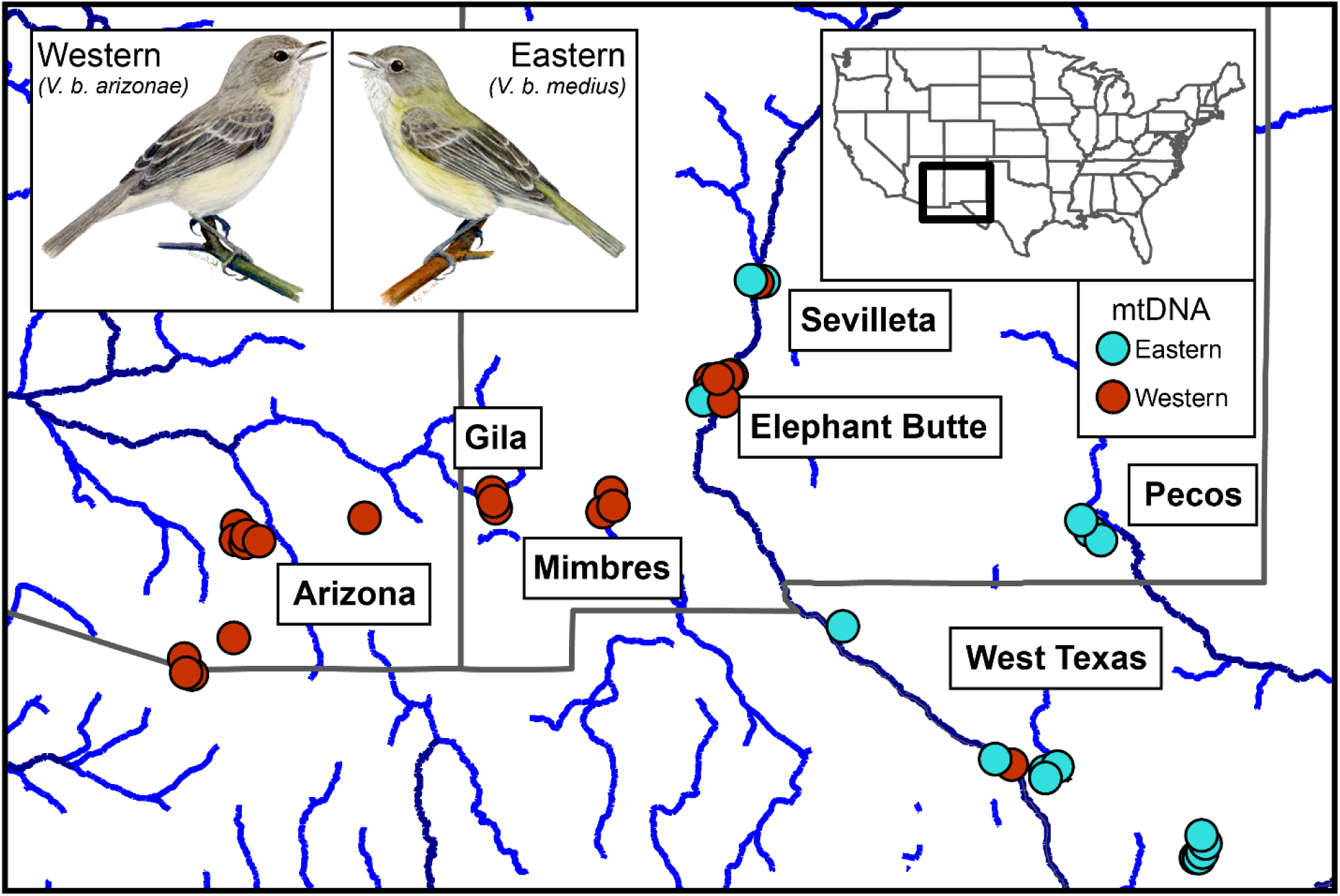
A) Map of the focal sampling region. Top left inset has depictions of study taxa, representatives of *Vireo bellii arizonae* and *V. b. medius*, the focal western and eastern populations, respectively (art by ABJ). Top right inset shows focal region of the United States. Dark gray lines represent state boundaries. Blue lines represent relatively major drainages, with the darker blue representing the higher flow ones. Sampling points are colored by the mitochondrial haplogroup of that individual, with boxes indicating sampling site names.

Genomic DNA was extracted using QIAgenDNeasy extraction kits (Qiagen Inc., Valencia, CA, USA) following manufacturer’s protocols. We quantified the amount of DNA using a Qubit 3.0 Fluorometer. One sample for use in 10X chromium sequencing was extracted at Discovery Life Sciences using a high molecular weight extraction for optimal reference quality. We selected individual MSB:Bird:60504 from the Gila River drainage because individuals from nearby sites in Arizona were thought not to be admixed based on a previous genetic study (Klicka et al. 2016).

### Sequencing

Two genomic sequencing methods were used: 10X Chromium sequencing for the reference genome assembly and whole-genome resequencing for all other samples. The reference genome was prepared and sequenced on 18% of an Illumina NovaSeq S4 lane by Discovery Life Sciences. Libraries for the remaining 43 samples were prepared at the University of New Mexico following the KAPA library preparation protocol and sequenced primarily on a single lane of a NovaSeq S4 at the Oklahoma Medical Research Facility. One additional sample was sequenced alongside other projects on a partial lane at the same facility. We selected three resequencing libraries representing individuals from the Sevilleta, Pecos, and Missouri for higher coverage sequencing to facilitate Pairwise Sequential Markovian Coalescent (PSMC) analyses, enabling assessment of effective population size variation across New Mexico relative to the core species range.

### Reference Genome

The reference genome was assembled using Supernova v2.1.1 (10X Genomics), optimized for Chromium sequencing data. Parameters were set for 56x coverage based on preliminary analyses; all other parameters were set to default. We then used ARKS v1.2.2 (Coombe et al. 2018) and LINKS v1.8.7 (Warren et al. 2015) to correct the assembly for downstream analyses. Due to the relatively fragmented nature of this genome, we performed reference-based scaffolding using RagTag v2.0.1 using the Hooded Crow (*Corvus cornix*) reference genome as a guide (Poelstra et al. 2014). We used QUAST v 5.0.2 (Gurevich et al. 2013) to calculate summary statistics about the reference and BUSCO v5.2.1 (Simão et al. 2015) to assess genome completeness as measured by the number of expected genes detected. This genome was used as input for population genomic analyses.

### Reference Annotation

To identify genes potentially underlying adaptation, we annotated first repeats and then genes on our scaffolded genome, largely following Eliason et al. (2022). First, we modeled repeats de novo using four iterations of RepeatModeler v2.0.5 (Flynn et al. 2020), combining these results with known Zebra finch (*Taeniopygia guttata*) repeats as input for RepeatMasker v4.1.5 (Smit et al. 2015). We performed structural annotation using a homology-based approach in GeMoMa v1.9 (Keilwagen et al. 2018). We used two high-quality passerine genome annotations for inference: Zebra Finch (Warren et al. 2010) and Hooded Crow (*Corvus cornix*; Poelstra et al. 2014).

### Variant Calling of Nuclear and Mitochondrial DNA

Sequences were aligned to the reference genome using BWA MEM v0.7.17 (Li & Durbin 2009). We performed two different methods of variant calling from whole genomic data, one for PSMC analysis and the other for all libraries used for population genomic analyses. For PSMC analysis, we used a pipeline of SAMtools v1.12 (Li et al. 2009), Picard (Broad Institute), and BCFtools v1.12 (Li et al. 2009) to call variants and generated a diploid consensus sequence as input for the analysis. This pipeline was run for the five individuals sequenced at high coverage and for the reference genome. For all 43 resequenced genomes, we followed a pipeline based on the Genome Analysis Toolkit’s (GATK; McKenna et al. 2010) Best Practices to call Single Nucleotide Variants for downstream population genomic analyses. This approach entailed marking duplicates and sorting the output of BWA-MEM using the GATK v4.1.9 tool MarkDuplicatesSpark, then calling and genotyping variants separately for each scaffold of the reference genome assembly. We did this using GNU Parallel (Tange 2018) and the GATK tools HaplotypeCaller, GenomicsDBImport, and GenotypeGVCFs. The resulting per-scaffold VCFs were combined using GATK’s GatherVcfs, then filtered with SelectVariants and VariantFiltration, keeping only single nucleotide polymorphisms with a depth of coverage greater than 2, quality score greater than 30.0, and quality-by-depth greater than 2. We further pruned these VCFs using VCFtools v0.1.15 (Danecek et al. 2011) to account for linkage disequilibrium.

We used a similar pipeline to that used for PSMC (SAMtools, Picard, and BCFtools) to generate mitochondrial DNA sequences for birds not sampled by Klicka et al. (2016). Unlike the pipeline used for PSMC, we did not permit heterozygous sites. We used a sequence of the ND2 gene from Klicka et al. (2016) as a reference (NCBI GenBank KM262750.1). We aligned reads from each whole-genome resequencing library to this single-gene reference and used those reads to determine if that mitochondrial haplotype was from the eastern or western haplotype group.

### Population Structure

We assessed population structure of *V. bellii* using three approaches. First, we used the glPca method within adegenet v2.1.10 (Jombart 2008) using a 75% complete VCF for our focal samples to conduct a principal component analysis to obtain a non-model-based estimate of population structure. Input was generated with vcfR v1.15 (Knaus & Grünwald 2016). Second, we estimated ancestry coefficients using ADMIXTURE v1.3.0 (Alexander & Lange 2011), with input generated using VCFtools v0.1.15 (Danecek et al. 2011) and PLINK v1.90b6.21 (Chang et al. 2015). We also calculated F_ST_, an index that assesses population divergence, using VCFtools. In addition to standard population structure analyses, we generated a phylogenetic network (Neighbor-net) with SplitsTree v4.14.2 (Hudson 1998) using input distances estimated by calculating Nei’s D with StAMPP v1.6.3 (Nei 1972, Pembleton et al. 2013). Note that this dataset had individuals with a mean depth less than 4x filtered.

To visualize population structure on the landscape, we used the program Estimated Effective Migration Surfaces (EEMS; Petkova et al. 2016). We generated input with PLINK v2.00a5.12 (Chang et al. 2015) and EEMS’s bed2diffs program. We then ran twenty replicate chains of EEMS with different seeds, 3.5 million MCMC iterations (discarding the first 500 thousand as burn-in), and 100 demes. We visualized MCMC chains to confirm convergence and plotted effective migration and diversity surfaces using reemsplot2. This analysis was repeated for a dataset of only autosomes and only the Z chromosome to account for the different patterns of course-scale population structure observed in them.

### Demography and diversity

We used three metrics to assess the effective population size and demography of *V. bellii* populations. First, we assessed per-site heterozygosity (what percentage of sites had two different alleles) of the individual used for the reference genome, as calculated during assembly with Supernova v2.1.1 (Weisenfeld et al. 2017). Second, we used an implementation of the PSMC (Li & Durbin 2011) to leverage our reference genome and genomes sequenced to a higher coverage to produce an estimate of effective population size throughout time (Allendorf et al. 2010, Oh et al. 2019). We also used pixy v1.2.3 (Korunes & Samuk 2021) to calculate population-level nucleotide diversity, using input that included invariant sites to avoid biased estimates of nucleotide diversity (Korunes & Samuk 2021). We calculated nucleotide diversity at both a subspecies level and for individual populations as defined by clusters of sampling sites.

### Genomic Landscape of Divergence

We visualized the genomic landscape of divergence using Weir & Cockerham’s weighted F_ST_ calculated in 10kbp and 50kbp windows using VCFtools (Danecek et al. 2011), which were plotted in qqman v0.1.8 (Turner 2018). Regions exhibiting elevated divergence were cross-referenced with gene annotations to identify potential functional relevance. In addition to this common approach, we aimed to determine how between and within population diversity contributed to regions of elevated F_ST_, windowed metrics for F_ST_, between population diversity (π_B_, also known as d_xy_), and within population diversity for a given pair (π_W_, with specific cases being referred to by their locality). Although the mean π_W_ for a pair is used for calculation of F_ST_, we looked at the effect that each population’s π_W_ has on the estimation of F_ST_ by taking a ratio of the two parameters. If uneven π_W_ is driving F_ST_ peaks, it would result in a positive correlation between this ratio and F_ST_.

### Simulations of Genomic Landscapes

To test the role asymmetric introgression plays in shaping the ratio of nucleotide diversity in areas of elevated F_ST_, we performed a series of population genetic simulations in SLiM v5.0.1 (Haller et al. 2025). These simulations did not attempt to replicate empirical parameters of the system at hand but rather test the role asymmetric introgression has in shaping the genomic landscape of divergence. Simulations followed a Wright-Fisher model and modeled three 100 Mbp chromosomes (two autosomes, one Z chromosome) with mutation and recombination rates of 2×10^-9^ per bp per generation. Each simulation started with a population of 10,000 individuals, which evolved for 50,000 generations before splitting into two populations of 10,000 individuals each for another 10,000 generations. After a total of 60,000 generations, new mutations were allowed to arise at a variable frequency (0.1% or 0.001% of new mutations) with a variable selection coefficient drawn from an exponential distribution with a mean of 0 (neutral) or 0.01 (beneficial). Finally, after 1,000 additional generations (61,000 total), bidirectional gene flow was permitted. Gene flow was permitted at either 0, 0.5, or 1 migrant per generation from population 1 to 2. From population 2 to 1, migrants per generation were permitted at either the same, half, or one tenth of that value. After gene flow began, samples of 50 individuals per population were taken in ten steps every 1,000 generations (10,000 total), and per-population diversity (as measured by π) and Wright’s F_ST_ were estimated in 200kb windows. This output was then read into R v4.4.1 (R Core Team 2024) for analysis and plotting.

## Results

### Sequencing

Whole genome resequencing was successful for all individuals, with some variance in individual coverage (Table S1). Samples targeted for lower coverage (n=38) ranged from 3.05– 15.8x mean depth, while higher coverage samples (n=5) ranged from 22.5–33.4x mean depth. The reference genome resulted in unexpectedly high coverage (∼70x) and was subsequently down-sampled to the recommended 56x.

### Reference Genome and Annotation

Our assembly, corrected with ARKS and LINKS, achieved a contig N50 132.24 kb and scaffold N50 of 7.31 Mb. The RagTag-scaffolded reference was chromosome-level; however, the chromosome arrangements likely differ from the true synteny in *V. bellii* due to using the Hooded Crow reference. Our homology-based annotation identified 15,541 genes with a mean length of 17.5 kbp. Of the two reference species used for annotation, the more closely related Hooded Crow contributed more annotated genes (13,464 genes, 7,236 unique) than the Zebra Finch (8,305 genes, 2,077 unique).

### Mitochondrial DNA

Analysis of mitochondrial DNA from individuals not included in previous studies (Klicka et al. 2016) yielded three key findings. First, individuals from far eastern and western New Mexico carried mitochondrial haplotypes consistent with populations from Texas and Arizona, respectively. Second, unexpectedly, individuals from the Rio Grande Valley harbored a mixture of eastern and western mitochondrial haplotypes. Specifically, at Sevilleta NWR, two individuals had eastern haplotypes and one had a western haplotype, while north of Elephant Butte Lake, one bird had an eastern haplotype and seven had western haplotypes (Figure 1). Despite the apparent differences in frequencies, this pattern was not statistically significant (Fisher’s Exact Test p>0.10). Lastly, we detected mitonuclear discordance in a West Texas individual possessing a western haplotype, whereas an Arizona bird from a previous study carried an eastern haplotype (Klicka et al. 2016).

### Population structure

Population structure analyses revealed a complex pattern of population divergence, partially inconsistent with historical subspecies hypotheses (Figure 2). First, populations from Arizona, and the Mimbres and Gila drainages in New Mexico clustered together, consistent with the subspecies *V. b. arizonae*. Second, birds from West Texas and the Pecos drainage in New Mexico formed another distinct cluster, aligning with *V. b. medius*. However, contrary to the past hypothesis that birds from the Rio Grande Valley in New Mexico were *V. b. medius* (Hubbard 1970), these samples were genetically intermediate, exhibiting variable admixture proportions of eastern and western ancestry (Figure 2). Additionally, phylogenetic network analysis also showed evidence for reticulation with the characteristic “blocky” appearance to branches connecting samples from the contact zone, reflecting admixture and genetic variability (Figure S1).

**Figure 2:**
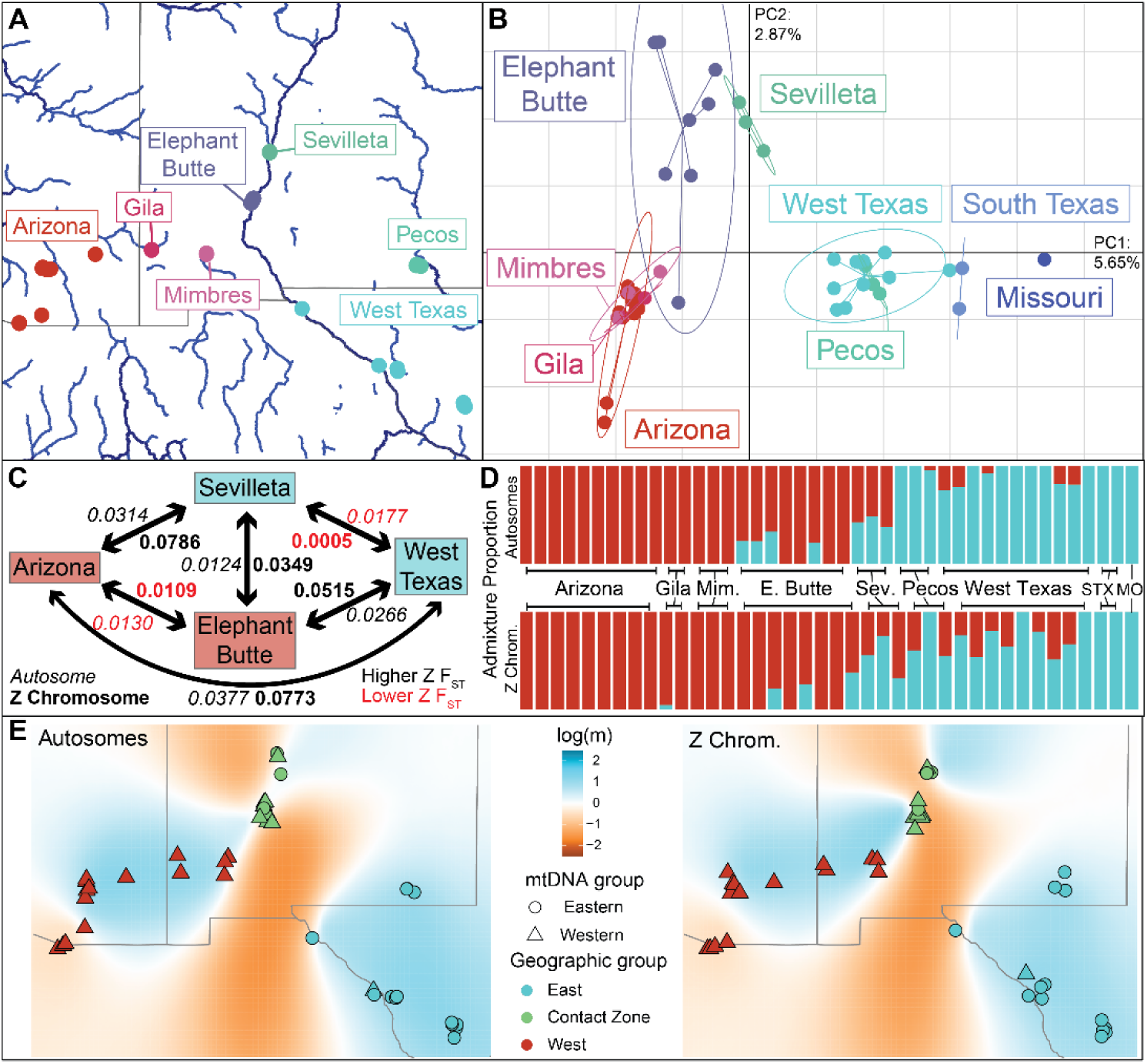
A) Map of the focal sampling region for reference, with points corresponding to sampling localities. B) Principal component analysis of genomic data for all samples, only including autosomes. C) Values of pairwise weighted Weir and Cockerham’s (1984) F_ST_ for four focal populations. Italicized values correspond to autosome-only estimates, and bolded values are Z chromosome only. Red text denotes comparisons where the Z chromosome value is lower than the autosomal value. D) Admixture analysis of focal samples for autosomes (top) and Z chromosome only, removing three outlier individuals from principal component analysis. Abbreviated populations in order are: Mim = Mimbres, Sev = Sevilleta, STX = South Texas (Lower Rio Grande), and MO = Missouri. E) Effective migration surfaces for autosomes (left) and the Z chromosome (right). Colors of points correspond to broad nuclear groups, and shape with mitochondrial group. The color of the map denotes areas of low gene flow (orange) and high gene flow (blue).

Despite being farther from core eastern populations in West Texas than Elephant Butte samples are, birds from the Rio Grande at Sevilleta NWR were genetically more similar to eastern populations compared to those from Elephant Butte. This suggests a source of western genetic ancestry concentrated around Elephant Butte. Effective migration surfaces (EEMS) analysis supported a narrow gene flow corridor connecting our sampling points along the Middle Rio Grande Valley (Figure 2E), but we note that the exact location and width of this corridor is likely influenced by exact sampling localities. However, it clearly reflects the relative lack of isolation compared to proximate parental populations, such as the Mimbres and El Paso sampling localities.

Although we inferred the New Mexico Rio Grande populations as a corridor of gene flow between eastern and western populations, we inferred it was a stronger barrier for sex-linked alleles than autosomal. This can be seen in the EEMS results limited to only the Z-chromosome, which inferred an overall similar pattern to autosomal DNA but with a barrier between Sevilleta and Elephant Butte samples (Figure 2E). This excess of divergence for the Z-chromosome was also by F_ST_ (Figure 2C), despite signs of admixture beyond the bounds of the Middle Rio Grande (Figure 2D). Although elevated divergence on the Z chromosome is expected, it is particularly high in contact zone populations (2.8 times autosomal, although still low at F_ST_=0.0349).

Although Z chromosome F_ST_ was substantially elevated for most comparisons, it did not hold true when comparing the Z-chromosome F_ST_ between Middle Rio Grande populations and adjacent ones with closer nuclear and mitochondrial affinities (Figure 2). Instead, the Z-chromosome F_ST_ was lower for those taxa, suggesting either a deficit of between-population diversity or excess of within-population diversity. Comparing the respective summary statistics to other populations, we found that the Elephant Butte population had significantly lower autosomal diversity relative to the Arizona population (Table S3, t-test p<<0.001), but non-significantly lower Z-chromosome diversity (p=0.097). This suggests a possible excess of diversity in the Elephant Butte-Arizona comparison. However, the pattern of relatively elevated Z-chromosome diversity was not recovered for the eastern-aligned population in a Sevilleta-Texas comparison, where diversity was significantly lower for the Sevilleta population genome-wide and on the Z-chromosome, and higher in magnitude overall (p<<0.001).

### Demography and diversity

The per-site heterozygosity, a proxy for population-wide nucleotide diversity, was 0.0045, which is intermediate relative to other passerines (Dutoit et al. 2016, Brüniche-Olsen et al. 2019). Results of our PSMC analysis of effective population size (Ne) through time suggested a stable (i.e., no more than a two-fold decline) population for the past million years for all sampled populations. However, the exact demographic pattern differed between populations (Figure 3). For western New Mexico populations, the population remained relatively stable during the Last Interglacial (Figure 3A), while eastern populations—including an admixed population with more eastern ancestry at Sevilleta NWR—increased (Figure 3B–D).

**Figure 3:**
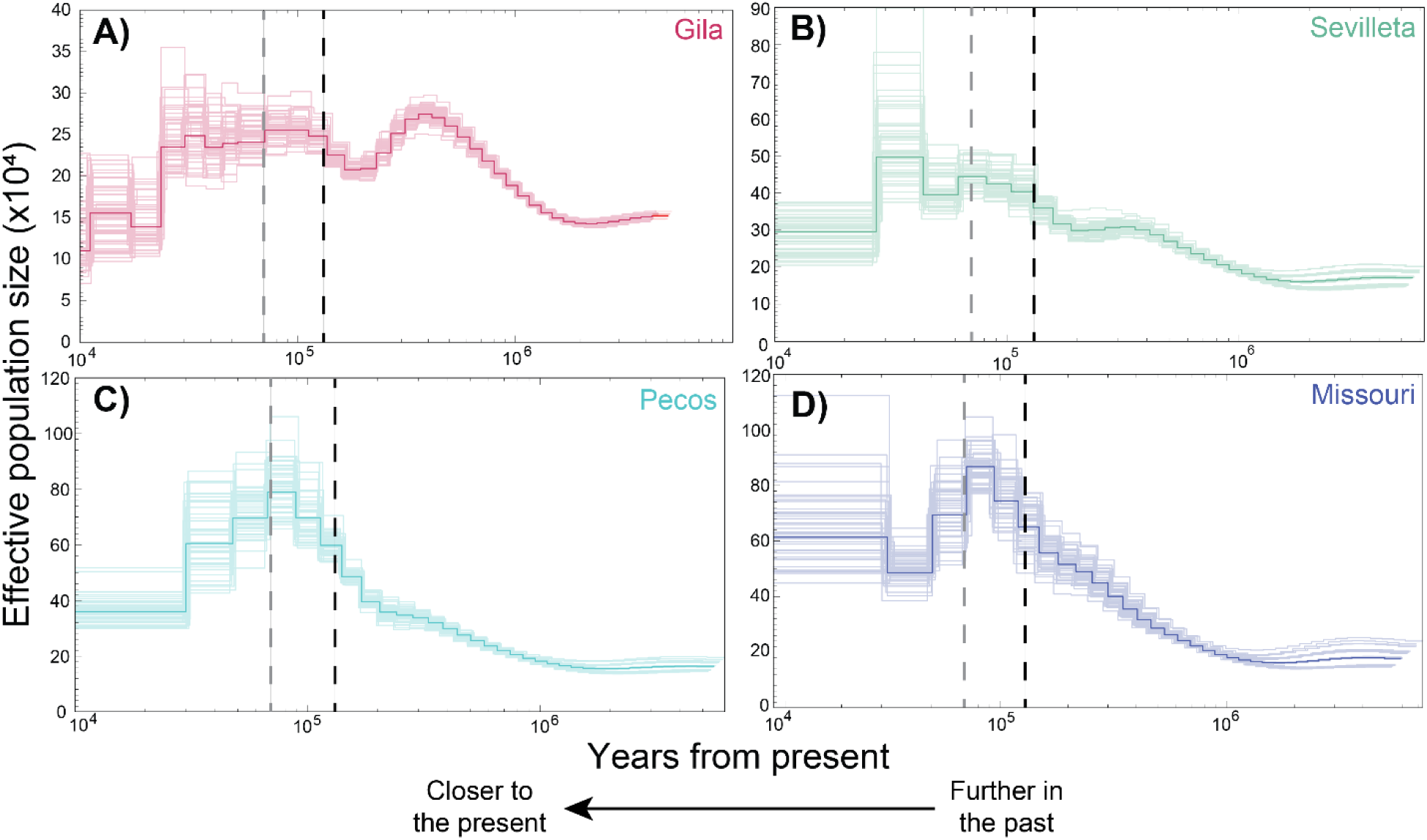
Plots of effective population size over time for four representative populations: the Gila river (A; western), Sevilleta NWR on the Rio Grande (B; admixed), the Pecos river (C; eastern), and Missouri (D; far eastern). Opaque lines are the absolute estimate, and translucent lines represent bootstraps. The black vertical dashed lines corresponds to the approximate time of the last interglacial (i.e., the last period of warmth before the last glacial period; approximately 120,000 years ago). The gray vertical lines represents the end of the mid-Pleistocene transition (approximately 70,000 years ago)), after which ice sheet extent increased dramatically (Willeit et al. 2019). Note that the y-axis differs for each sub-plot while the x-axis is consistent across plots.

Population-level estimates of nucleotide diversity (π) performed with pixy v1.2.3 (Korunes & Samuk 2021) were consistent with estimates from whole genome data and PSMC analyses (Table S3). Western populations (namely Arizona) had lower nucleotide diversity (especially on the Z chromosome), while eastern ones (West Texas) had higher diversity. The admixed populations displayed reduced diversity relative to both, likely due to occupying a small geographic area. This could explain their clustering apart from other populations on the second principal component of our PCA (Figure 2B).

### Genomic landscape of divergence

Standard analyses of the genomic landscape of divergence found a pattern familiar to those of many non-model organisms: a baseline low-level of F_ST_ punctuated by few peaks that include much of the sex-linked Z chromosome (Figure 4A). However, only a few unique regions exceeded the 99.95^th^ percentile of windowed F_ST_, our threshold for notability. Two other large regions surpassed this threshold in our focal comparisons: a 1Mb peak on Chromosome 1 in the Texas–Arizona comparison (Figure 4, first column) and a 1.1 Mb peak on Chromosome 12 in the Sevilleta–Elephant Butte comparison (Figure 4, third column).

**Figure 4:**
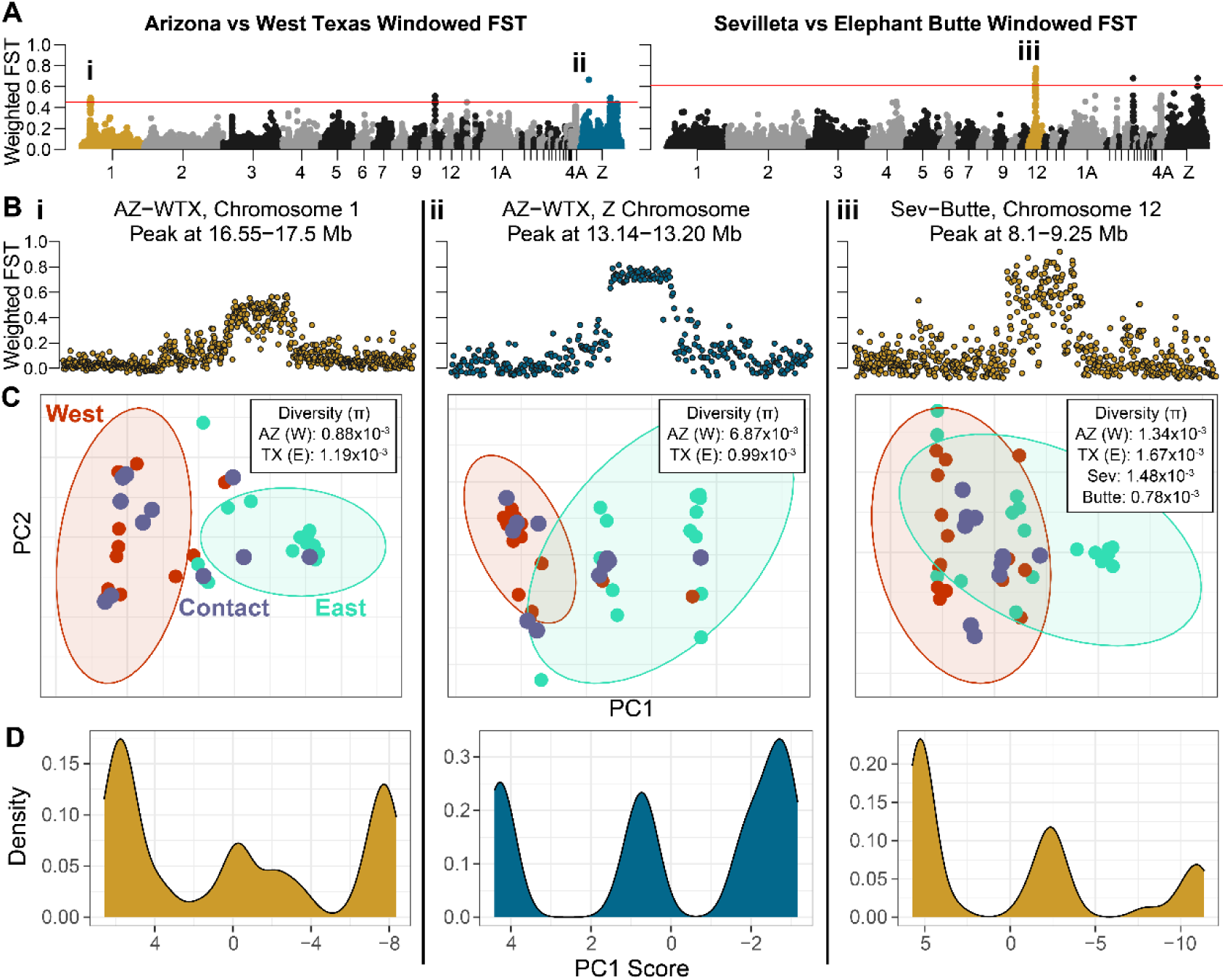
Plots of the genome landscape of divergence, highlighting three especially strong and well defined outlier regions in both of our focal F_ST_ comparisons: Arizona vs West Texas and Sevilleta vs Elephant Butte. A) Manhattan plots of weighted Weir & Cockerham’s estimator of F_ST_ calculated in 50 kb windows across the whole genome, with peaks labeled to correspond to the rest of the figure. Note that ii is only the single small peak, not the lower, wider one. Red line denotes windows at or above the 99.95^th^ percentile. B) Zoomed in Manhattan plots for the three regions in 10 kb (i and iii) and 1kb (ii) windows, highlighting the consistent height and strong demarcations. For ii, the x-axis scale is lower (peak is ∼40 kb, others are ∼1 Mb). Non-peak values are displayed, but range refers only to the peak. C) Principal component analysis of variants in these peaks. Colors of point and ellipses represent sampling region, with West (red) corresponding to Arizona, Gila, and Mimbres; Contact (purple) Sevilleta and Elephant Butte; and East (teal) corresponding to Pecos, West Texas, South Texas, and Missouri. Inset boxes represent nucleotide diversity for subsets of those populations, with TX corresponding only to West Texas. D) Kernel density plots of PC1 values for the plots in C, note weaker trimodality in i (left column) than in ii and iii (center and right columns, respectively).

These peaks were characterized by reduced within-population diversity, but one region on the Z chromosome stood out for also exhibiting elevated between-population divergence in the Texas-Arizona comparison (Figure S2). This region showed both sharply defined peaks in the Manhattan plot and a strongly tri-modal pattern in the principal component analysis, consistent with a small (61 kb) inversion (Figure 4, second column). Although the tri-modal PCA pattern did not show a significant deficit of heterozygotes (HWE Chi-sq p-value > 0.05), the region appeared nearly fixed in the western populations and highly variable in the eastern population. Gene annotation identified only one gene in this region: 3-oxoacid CoA-transferase 1 (OXCT1), which encodes a mitochondrial matrix enzyme involved in ketone metabolism (Ozsvari et al. 2017). The two larger peaks also showed sharply defined peaks, with either weakly (Figure 4, first column) or strongly (Figure 4, third column) tri-modal PCAs. These regions contained many more genes (29 and 27, respectively), none of which were clearly related to mitonuclear function in our preliminary survey.

Our exploration of population-specific contributions to the landscape of divergence revealed a pattern similar to that observed in the putative inversion, with divergence peaks largely driven by reduced diversity in Arizona population (Figure 5A). To be precise, the ratio between the windowed nucleotide diversity in the putative sink (West Texas) and putative source (Arizona) is highest in the most divergent windows. These results suggest that the overall landscape of divergence is shaped primarily by a reduction in genetic diversity in western (Arizona) populations—a conclusion supported by simple linear models. For example, in the Texas-Arizona comparison, adjusted R^2^ values for individual components of F_ST_ (i.e., within-population and between-population diversity) ranged from 0.091 to 0.138. The R^2^ value for their ratio—which does not directly contribute to the calculation—was 0.336. This pattern was consistent across multiple sets of comparisons but was elevated on the Z chromosome. Two of our three focal peaks also reflected this pattern: the small Z-linked region (ii) represented the most extreme value in Figure 5A for the Texas-Arizona comparison, while the Chromosome 12 peak (iii) accounted for much of the elevated diversity ratio in the Sevilleta-Elephant Butte comparison.

**Figure 5:**
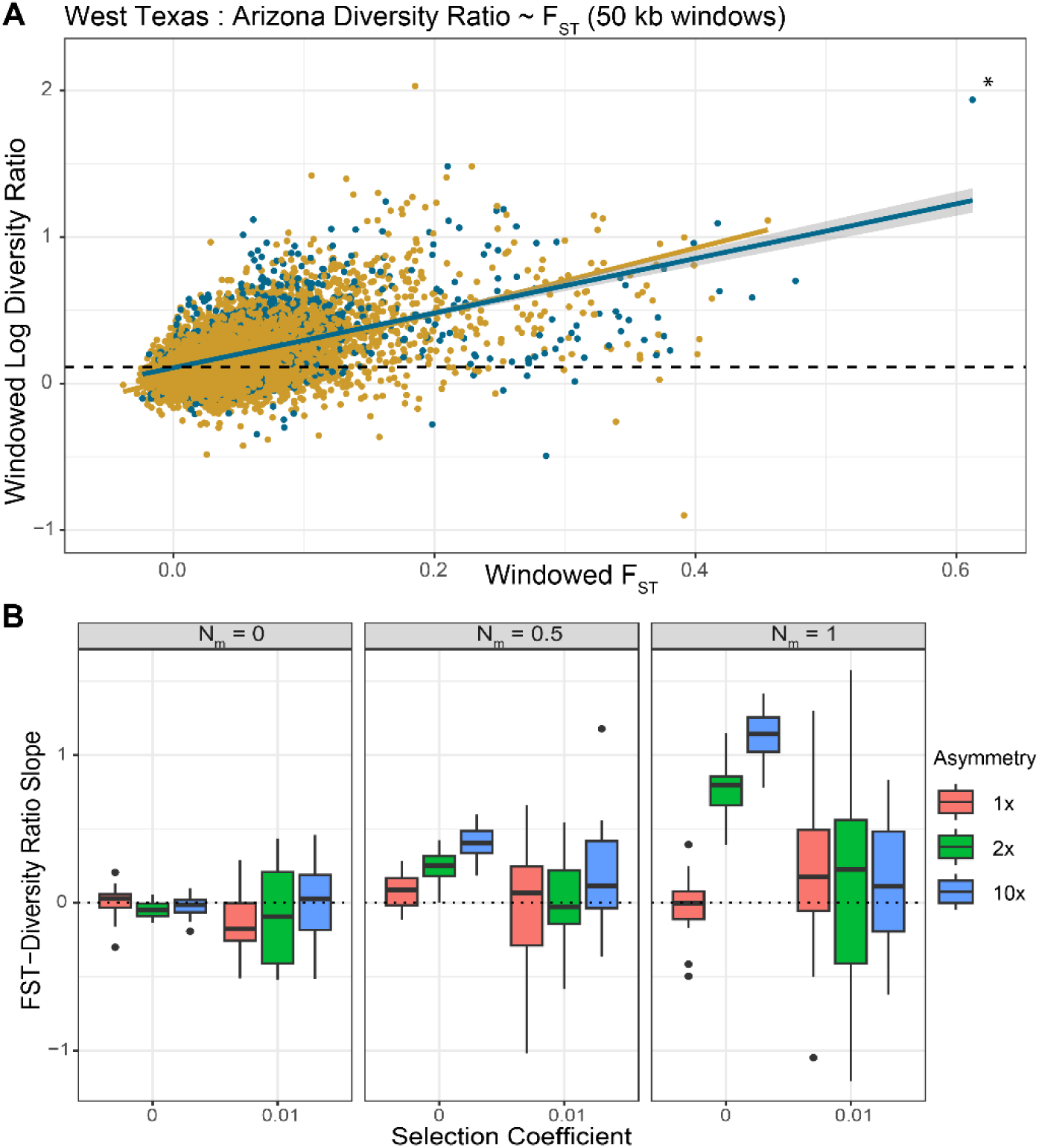
A) Plot exploring how population-specific nucleotide diversity drives elevated F_ST_. Each displays the natural log of the ratio of nucleotide diversity for the eastern (West Texas) to the western (Arizona) population. Higher positive values indicate a higher nucleotide diversity in the first population, meaning that the elevated F_ST_ (for high values) is driven by reduced diversity in the second population. All statistics calculated by pixy in 50 kb windows. Both trend lines are highly significant (p<<0.001). Starred point in top left represents the putative Z-13M inversion (Figure 4). B) Simulations of the regression coefficient for the line above, with the ratio being between the net sink of migrants and the net source of migrants. Positive values denote that higher F_ST_ values are driven by reduced variation in the source population, and negative values the sink population. Note that the slope of the regression of all windows in A is 2.23 (i.e., greater than simulation values). The three plots have varying base levels of migration (N_m_=0, 0.5, and 1), the x axis denotes simulations with (right) and without (left) positive selection, and the fill denotes the degree of asymmetry (e.g., 10x with N_m_ = 1 means the source receives 0.1 migrants/generation).

### Simulating the genomic landscape of divergence

Our population genetic simulations revealed that the observed divergence pattern is consistent with asymmetric introgression of neutral loci, which consistently yielded a positive slope (Figure 5B). However, despite the strong trend, the simulated slopes were weaker than in empirical data (2.23 empirically, mean of 1.13 for the highest level of asymmetric gene flow in Figure 5B). Additionally, consistent with the eastern empirical data, the mean genome-wide diversity was higher in the sink population (diversity ratio of approximately 1.13 empirically, 1.4 in simulations with highest mean slope in Figure 5B). In short, regions with elevated F_ST_ were primarily driven by reduced diversity in the net source population. While simulations incorporating positive selection also produced notable slopes, the results were far more variable—likely due to the relatively strong selection coefficients used, which generated a mixture of partial and complete selective sweeps.

## Discussion

We demonstrated a unique and unexpected pattern of divergence in New Mexico’s *Vireo bellii* populations. First, samples from western populations in New Mexico and Arizona showed minimal evidence of genome-wide admixture while samples from West Texas and the Pecos drainage showed some signs of more western ancestry (Figure 2). However, the strongest signs of admixture were in birds along the Rio Grande in central New Mexico. These populations formed a discontinuous contact zone, with an excess of western ancestry on the part of the Rio Grande (Elephant Butte) adjacent to the mountainous Gila region of southwest New Mexico while individuals at Sevilleta NWR showed a more even contribution from eastern and western populations. This is surprising, as the Sevilleta samples are more geographically distant from the core range of *V. b. medius* in West Texas than the Elephant Butte samples. This suggests that the isolated, north-south oriented habitat corridor presents an unusual biogeographic setting that we hypothesize enabled the formation of this discontinuous pattern of population structure. In short, we believe an initial colonization of the eastern population following the last glacial period, then asymmetric west-to-east gene flow facilitated by the fragmented drainages east of the Black Range—which contains the Continental Divide and overlooks the Rio Grande—resulted in a geographically limited but impactful region of gene flow (Figure 6).

**Figure 6:**
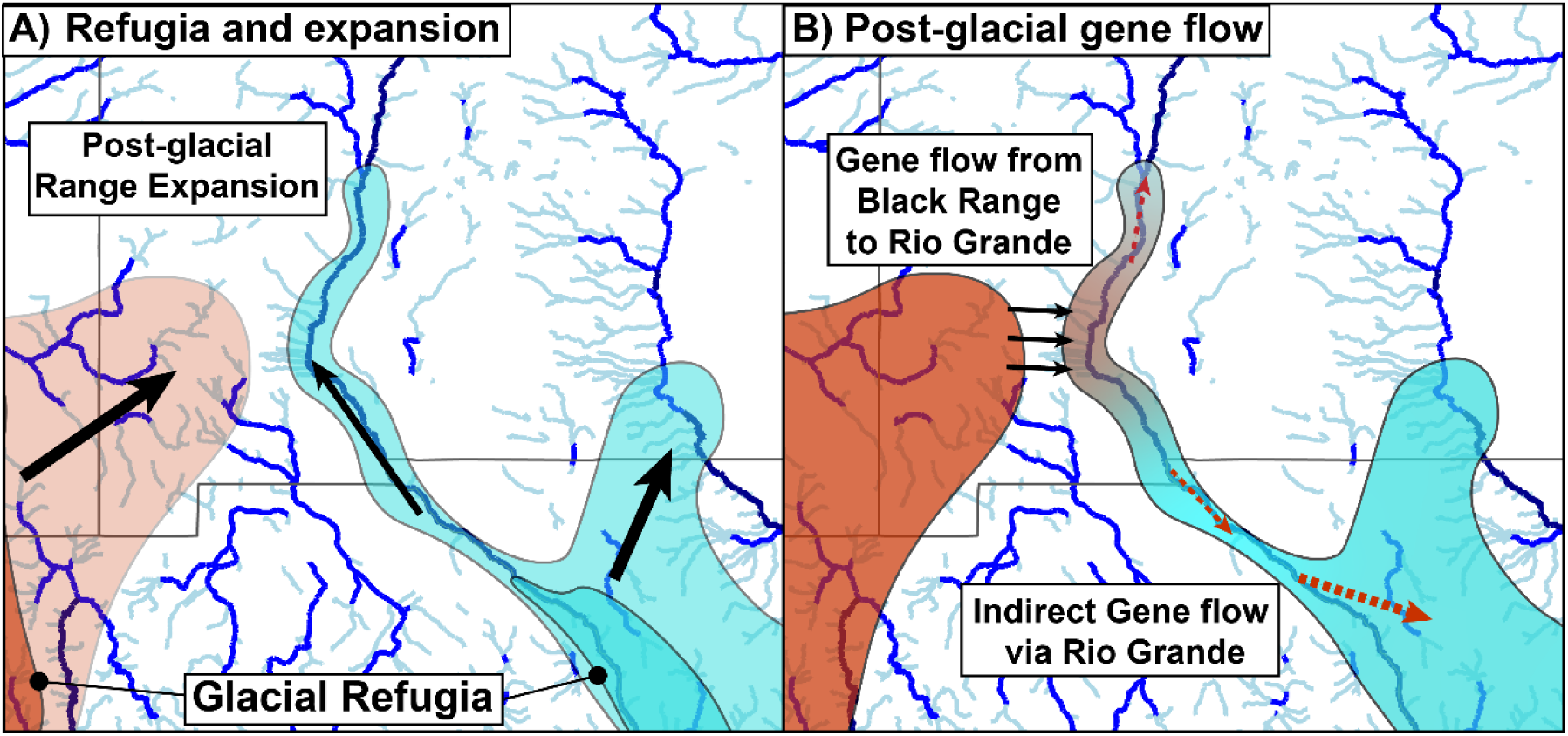
A map outlining our hypothesized history of expansion and gene flow in *Vireo bellii*. A) Depicts approximate glacial refugia (based on Klicka et al. 2016) in dark, and pale representing expansion to the current range, notably with initial colonization of the Rio Grande from the eastern population. B) Depicts hypothesized current patterns of gene flow, with direct gene flow to the region around Elephant Butte via drainages coming from the Black Range and Mimbres drainages (black arrows), and indirect gene flow along the Rio Grande to both the Sevilleta and Texas.

We also found evidence that the same asymmetric gene flow driving the contact zone shaped patterns of genome-wide divergence. While the overall genomic landscape resembles that of other recently diverged avian taxa (Figure 4A; Irwin et al. 2018), we show that these differences are driven in part by large, highly divergent regions—consistent with chromosomal inversions—which harbor markedly reduced diversity in western populations compared to eastern ones (Figure 4B-D). This asymmetry in diversity was recovered across the genome under regions of elevated divergence (Figure 5A), and our simulations demonstrate that asymmetric introgression alone, even in the absence of selection, can produce this pattern (Figure 5B).

### Biogeographic history of *Vireo bellii*

The two sampled regions along the Rio Grande in New Mexico revealed an unexpected pattern: the more northern population (Sevilleta NWR) showed higher *V. b. medius* ancestry than the population farther south. This result could suggest that the Rio Grande corridor was initially colonized by *V. b. medius,* or by an admixed population with a greater proportion of *V. b. medius* ancestry, moving northward; subsequent low-level gene flow from nearby *V. b. arizonae*, just to the west, altered the genetic makeup in one portion of the corridor (Figure 6). Supporting this interpretation, western Texas populations also showed some admixture with western (*arizonae*) birds (Figure 2D). This is consistent with genome-wide F_ST_ values, which finds that individuals from western Texas are much less divergent from *V. b. arizonae* (West Texas-Arizona F_ST_=0.038) than are individuals from southern Texas (South Texas-Arizona F_ST_=0.075; Table 1). This gene flow could have been facilitated by short-jump dispersal from the Gila River drainages (west of the Continental Divide) over mountain passes into fragmented riparian habitats such as the Mimbres River and Animas Creek on the east side of the Continental Divide, in the Rio Grande watershed.

Our findings align with past niche modeling (Klicka et al. 2016), which provided a framework for understanding the evolutionary history of *V. bellii* over the past million years. Eastern and western lineages followed distinct population trajectories: eastern populations expanded during the mid-Pleistocene, whereas western populations remained small and stable—reflecting both their historical and present-day range limitations. The eastern population’s growth began just before the end of the last interglacial (approximately 130,000 years ago; black dashed line, Figure 3) and continued through the mid-Pleistocene transition (∼70,000 years ago; gray dashed line, Figure 3). The eastern population then declined from the mid-Pleistocene transition to the present, likely driven by increasing glaciation and associated range contractions (Klicka et al. 2016). After the last glacial period, ranges expansions brought eastern and western populations into secondary contact along the Rio Grande in New Mexico, producing the current pattern of admixture observed today (Klicka et al. 2016).

### Taxon-specific implications

Although *Vireo bellii* is considered a threatened species in New Mexico due to its limited riparian habitat, our findings suggest relatively good genetic health across populations. Effective population sizes were consistently high (all >100,000, Figure 4a–c), and genetic diversity was moderate when compared to genome-wide estimates from other passerines (Table 2; Dutoit et al. 2016, Brüniche-Olsen et al. 2019). These results indicate that *V. bellii* is not currently at risk of losing genetic diversity due to drift. Nonetheless, continued habitat loss along the Rio Grande warrants extreme concern, as these central NM populations may be critical for maintaining range-wide genetic variation. A similar role has been noted for the contact zone between *V. b. arizonae* and *V. b. pusillus* in the Mojave Desert, where *V. b. arizonae* also acted as a net source of migrants. (Vandergast et al. 2024). In both cases, continued gene flow depends on the continued ecological integrity of small, intermittent aridland streams.

Empirical and simulation studies (Robinson et al. 2019; Kyriazis et al. 2021) further suggest that populations transitioning from historically large to small sizes can result in the expression of deleterious, recessive alleles, thereby increasing extinction risk. This highlights the potential vulnerability of the isolated riparian populations in central New Mexico. Gene flow across such contact zones can mitigate these risks by promoting adaptation to changing climatic conditions and facilitating genetic rescue from deleterious alleles (Brown & Kodric-Brown 1977, Oziolor et al. 2019).

From a taxonomic perspective, our data support the presence of two distinct lineages of *Vireo bellii* in New Mexico*—V. b. arizonae* and *V. b. medius*—that come into contact along the Rio Grande with some degree of gene flow. However, reproductive isolation appears weak or absent, as evidenced by admixed populations along the Rio Grande and gene flow extending into West Texas. The sharp genetic turnover observed between *V. b. arizonae* populations north of Elephant Butte Lake and *V. b. medius* along the Rio Grande in West Texas is especially striking, given the lack of obvious dispersal barriers within this shared habitat corridor. We hypothesize that this pattern reflects historical processes—specifically, range expansions following the last glacial maximum—rather than a stable zone that is maintained by reproductive incompatibility (Figure 6).

Regarding the subspecies status of New Mexico populations, our data indicate that birds in the Pecos drainage align with *V. b. medius*, while those from the Gila and Mimbres drainages represent *V. b. arizonae*. Delimiting hybrid populations is a taxonomic challenge (McCullough et al. 2021), so we recommend treating the Rio Grande populations as admixed rather than attempting formal subspecies assignment. Although we did not make note of behavioral characteristics that differentiate populations (e.g., tail wagging; Greaves et al. 2006), a preliminary examination of plumage revealed individuals from Sevilleta NWR were generally brighter than those from near Elephant Butte Lake consistent with their higher eastern ancestry proportion. Importantly, the geographic position of this contact zone makes it a likely conduit for maintaining range-wide genetic connectivity between divergent *V. bellii* lineages. Accordingly, as riparian habitats in the Southwest continue to contract, preserving these populations will be critical for long-term resilience of the species.

### Genomics of divergence followed by asymmetric gene flow

Our analysis of the genomic landscape of divergence yielded a surprising result: F_ST_ peaks were largely driven by differences in nucleotide diversity between the two best-sampled populations. At first glance, this pattern aligns with a model of secondary contact in which beneficial alleles sweep to fixation in the western population and then spread—partially or fully— through the eastern population. This is consistent with the “recurrent sweep” model of F_ST_ peaks (Cruickshank & Hahn 2014, Bay & Ruegg 2017, Irwin et al. 2018). However, we found that similar patterns could arise even in the absence of selection—through asymmetric gene flow alone (Figure 5B). Although this finding may seem counterintuitive, it reflects the fundamental principle that elevated F_ST_ is often driven more by reduced diversity than by divergent selection. When gene flow is asymmetric, introgression introduces new variation into the recipient population, increasing its diversity, while the source remains genetically reduced—thus elevating F_ST_.

This interpretation is supported by our population structure results, which show a disproportionate influx of western ancestry into the Rio Grande population north of Elephant Butte in central New Mexico. Following predictions from island biogeographic theory, the Rio Grande acts as a receptive “sink,” receiving migrants from larger western populations in the Mimbres and Gila drainages. These western populations likely serve as sources of migrants—and potentially beneficial alleles—that introgress into eastern *V. bellii*. This movement is likely facilitated by smaller seasonal drainages that connect the Black Range and the Gila region eastward to the Rio Grande (Figure 6). This might have included drainages such as Animas or Percha Creek, where Bell’s Vireos have been recorded during the breeding season (eBird; Sullivan et al. 2009). Populations as far as West Texas now show signs of genome-wide admixture, which we hypothesize occurred via this series of riparian corridors (Figure 2D).

While our results suggest that neutral processes—particularly asymmetric introgression—can fully explain the observed genomic patterns, we cannot rule out a role for selection. Indeed, our simulations incorporating selection did sometimes generate a positive relationship, despite the average being close to zero (Figure 5B). However, those simulations used relatively small population sizes and few genomic regions, and we believe that the central tendency is likely a better reflection of what a more complete simulation would recover. Future work with finer-scale sampling of Rio Grande populations could use a broader range of admixture fractions to test whether specific genomic regions under F_ST_ peaks are introgressing preferentially from the western to eastern background.

One plausible hypothesis for selection’s role in this system involves mitonuclear incompatibility, given the deep divergence between eastern and western mitochondrial haplogroups (∼1.11–2.04 Ma; Klicka et al. 2016). The disproportionately high mitochondrial divergence relative to nuclear divergence could reflect co-evolution between the two interacting genomes (Wang et al. 2021). In our dataset, the most divergent nuclear region (by absolute divergence) between eastern and western populations—apart from the mitochondrial genome itself—was a small inversion on the Z chromosome that contains *OXCT1*, a gene encoding a mitochondrial matrix enzyme (Ozsvari et al. 2017). This region stood out not only for its divergence but also for its distribution among populations: while it showed reduced diversity in western populations, it was widespread in eastern populations. Despite most individuals in West Texas and the Pecos River region carrying eastern mitochondrial haplotypes, we found a mix of heterozygotes and even homozygotes for the “western” inversion haplotype (Figure 4C, middle column). Although one could interpret this ratio as the results of adaptive mitonuclear co-evolution, alternative explanations—such as reduced recombination in the inversion maintaining divergence—should be considered.

Although we did not find a clear tie between this small inversion and reproductive isolation, we found some evidence for potential reproductive isolation associated with the Z chromosome. Across admixed populations, individuals showed reduced divergence on the Z chromosome from the more strongly represented parental lineage (Figure 2C), and patterns of inferred breaks in gene flow were strongest on the Z chromosome (Figure 2E). These patterns loosely mirror mitochondrial haplogroup frequencies, but our modest sample sizes at the Sevilleta NWR to date limit our ability to make firm conclusions. Nonetheless, this observation highlights a promising direction for future research. Denser sampling and targeted analyses could reveal whether specific regions of the Z chromosome are associated with eastern or western mitochondrial haplotypes and contribute to the observed pattern of divergence.

## Data Availability

Data will be made publicly available upon publication. Reads are uploaded to NCBI’s Short Read Archive (PRJNA XXX, in the process of uploading as of submission). All scripts and relevant files can be found on Dryad upon publication (https://doi.org/10.5061/dryad.z8w9ghxtm), and scripts and a subset of files can be found on GitHub (https://github.com/ethangyllenhaal/BellsVireoContact/).

## Acknowledgements

This work was primarily funded by the New Mexico Department of Game and Fish’s Share With Wildlife Grant. We thank the museums and staff that contributed non-New Mexico samples for this project, namely the San Diego State University Biodiversity Museum, Louisiana State University Museum of Natural Science (Donna Dittmann), University of Kansas Biodiversity Institute (Lucas DeCicco, Mark Robbins), and University of Texas El Paso Biodiversity Collection (Vicky Zhuang, Michael Harvey). We additionally thank Oscar Johnson, Vicente Mata-Silva, John Sproul, Indio Mountains Research Station, and Friends of the Rio Bosque for assistance obtaining certain West Texas samples. EFG was supported by National Science Foundation grants DGE-1650114 and DBI-2410565 while conducting this work. MJA was supported by National Science Foundation grant DEB-1557051. Field work conducted by LBK was funded by the Frank M. Chapman Memorial Fund of the American Museum of Natural History, American Ornithologists’ Union Research Award, and Los Angeles Audubon Society’s Ralph W. Schreiber Ornithology Research Award. We also thank Lisa Barrow, Benjamin Haller, Jeffery Long, Joseph Manthey, and Shannon Mindeman for assistance. We would like to thank the UNM Center for Advanced Research Computing, supported in part by the National Science Foundation, and the High Performance Computing Center at Texas Tech University for providing the high performance computing resources used in this work.

