## Supplementary material for "Asymmetric gene flow across a desert contact zone in a riparian songbird": Figure S1; Figure S2; Table S1; Table S2; Table S3

**Supplemental Information**

**Figure S1:** Phylogenetic network of sampled individuals, with larger labels denoting region of samples and smaller text at tips denoting sample names. Note the relatively high amount of reticulation at the New Mexico Rio Grande (i.e., Sevilleta and Elephant Butte) populations. Long branch lengths likely reflect the high genetic diversity of this species.

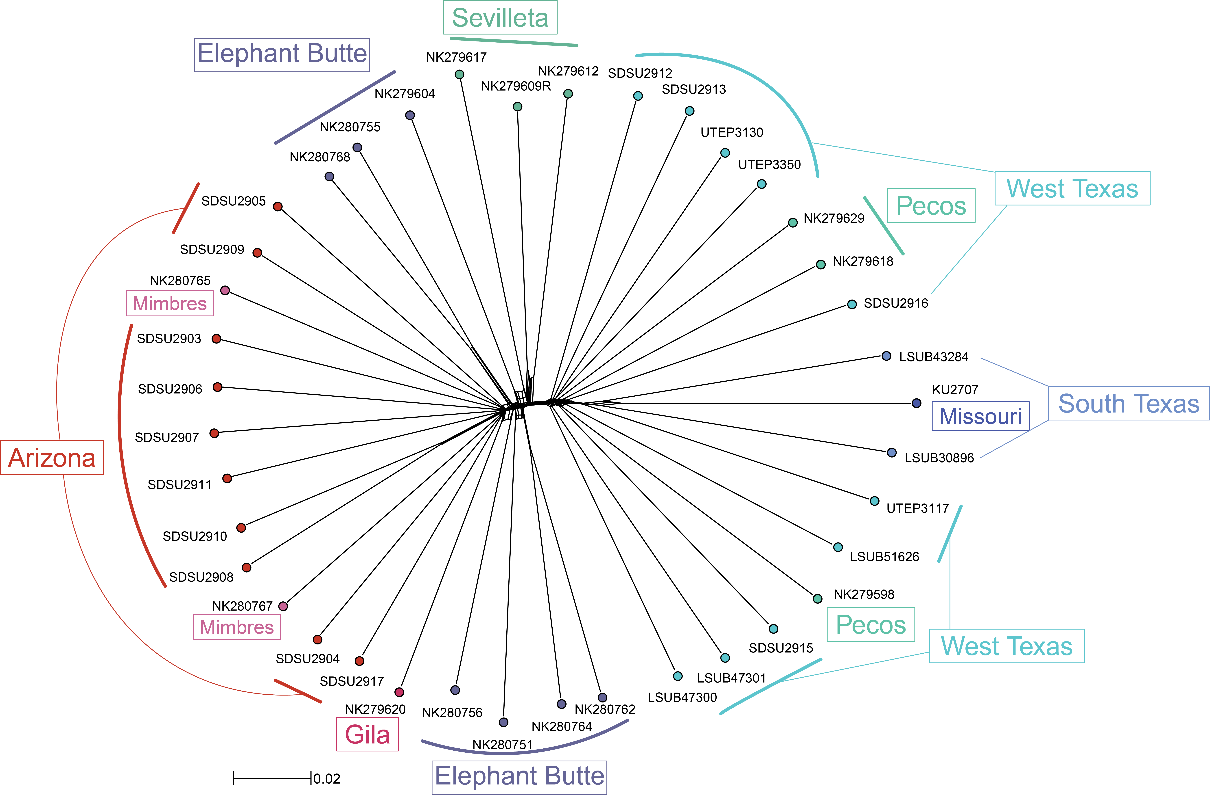

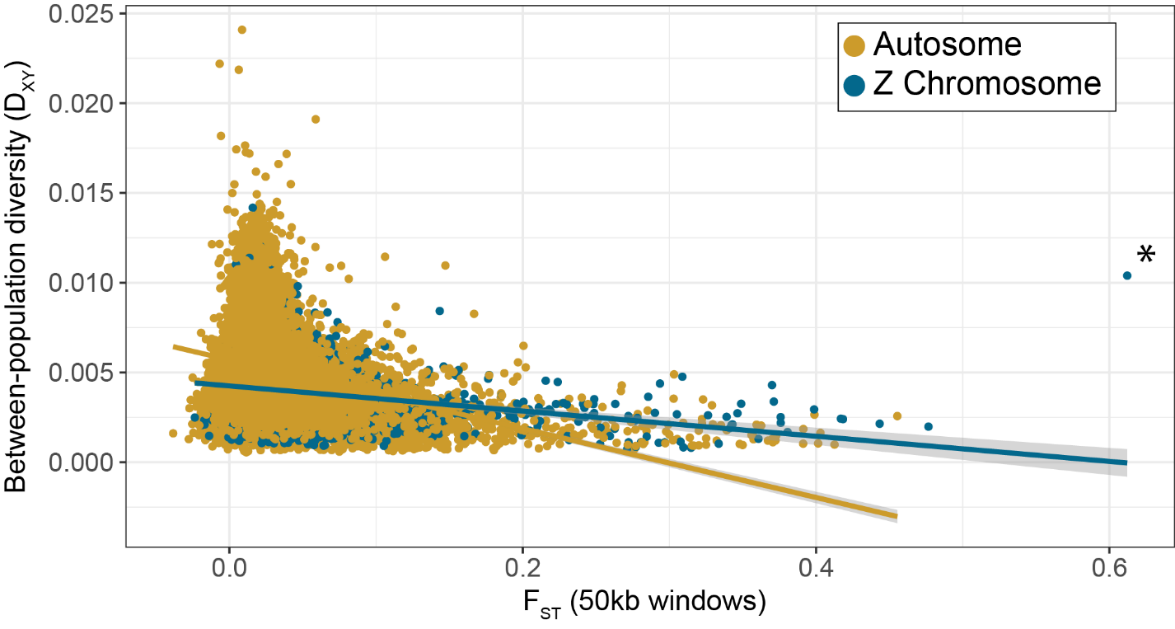
**Figure S2:** Plot of F_ST_ vs between-population diversity, calculated in 50kb windows in pixy. Trend lines are simple linear models plotted in R. The asterisk denotes the small Z outlier region.

**Table S1:** Sampling table for all samples included in this work. The first three columns are taxonomic identities (subspecies left blank for admixed populations), the next three are the institution and up to two identifiers, followed by the locality’s state and sampling region, then mitochondrial haplogroup and nuclear group, and finally depth of coverage. Bolded sample represents our reference genome.

| Genus | Sp. | Subsp. | Institution | Tissue | Voucher | Script name | State | Location | mtDNA | Nuclear | Depth | Arctos link |
| --- | --- | --- | --- | --- | --- | --- | --- | --- | --- | --- | --- | --- |
| Vireo | bellii | arizonae | SDSU | n/a | 2903 | Vireo_bellii_SDSU2903 | AZ | SE Arizona | West | AZ | 7.52 | n/a |
| Vireo | bellii | arizonae | SDSU | n/a | 2904 | Vireo_bellii_SDSU2904 | AZ | SE Arizona | West | AZ | 8.35 | n/a |
| Vireo | bellii | arizonae | SDSU | n/a | 2905 | Vireo_bellii_SDSU2905 | AZ | SE Arizona | West | AZ | 5.73 | n/a |
| Vireo | bellii | arizonae | SDSU | n/a | 2906 | Vireo_bellii_SDSU2906 | AZ | SE Arizona | West | AZ | 11.65 | n/a |
| Vireo | bellii | arizonae | SDSU | n/a | 2907 | Vireo_bellii_SDSU2907 | AZ | SE Arizona | West | AZ | 11.75 | n/a |
| Vireo | bellii | arizonae | SDSU | n/a | 2908 | Vireo_bellii_SDSU2908 | AZ | SE Arizona | West | AZ | 8.06 | n/a |
| Vireo | bellii | arizonae | SDSU | n/a | 2909 | Vireo_bellii_SDSU2909 | AZ | SE Arizona | West | AZ | 8.45 | n/a |
| Vireo | bellii | arizonae | SDSU | n/a | 2910 | Vireo_bellii_SDSU2910 | AZ | SE Arizona | West | AZ | 11.30 | n/a |
| Vireo | bellii | arizonae | SDSU | n/a | 2911 | Vireo_bellii_SDSU2911 | AZ | SE Arizona | West | AZ | 13.15 | n/a |
| Vireo | bellii | arizonae | SDSU | n/a | 2917 | Vireo_bellii_SDSU2917 | AZ | SE Arizona | West | AZ | 10.47 | n/a |
| Vireo | bellii | arizonae | MSB | 279620 | 60503 | Vireo_bellii_NK279620 | NM | Gila River | West | Gila | 8.43 | https://arctos.database.museum/guid/MSB:Bird:60503 |
| **Vireo** | **bellii** | **arizonae** | **MSB** | **279619** | **60504** | **Vireo_bellii_NK279619** | **NM** | **Gila River** | **West** | **Gila** | **Ref** | **https://arctos.database.museum/guid/MSB:Bird:60504** |
| Vireo | bellii | arizonae | MSB | 279607 | 60505 | Vireo_bellii_NK279607 | NM | Gila River | West | Gila | 3.80 | https://arctos.database.museum/guid/MSB:Bird:60505 |
| Vireo | bellii | arizonae | MSB | 280765 | 60557 | Vireo_bellii_NK280765 | NM | Mimbres River | West | Mimbres | 6.49 | https://arctos.database.museum/guid/MSB:Bird:60557 |
| Vireo | bellii | arizonae | MSB | 280767 | 60559 | Vireo_bellii_NK280767 | NM | Mimbres River | West | Mimbres | 7.57 | https://arctos.database.museum/guid/MSB:Bird:60559 |
| Vireo | bellii | arizonae | MSB | 280772 | 60564 | Vireo_bellii_NK280772 | NM | Mimbres River | West | Mimbres | 3.10 | https://arctos.database.museum/guid/MSB:Bird:60564 |
| Vireo | bellii | [admixed] | MSB | 279604 | 60501 | Vireo_bellii_NK279604 | NM | Rio Grande | West | Butte | 6.07 | https://arctos.database.museum/guid/MSB:Bird:60501 |
| Vireo | bellii | [admixed] | MSB | 280751 | 60543 | Vireo_bellii_NK280751 | NM | Rio Grande | West | Butte | 4.87 | https://arctos.database.museum/guid/MSB:Bird:60543 |
| Vireo | bellii | [admixed] | MSB | 280755 | 60547 | Vireo_bellii_NK280755 | NM | Rio Grande | West | Butte | 8.02 | https://arctos.database.museum/guid/MSB:Bird:60547 |
| Vireo | bellii | [admixed] | MSB | 280756 | 60548 | Vireo_bellii_NK280756 | NM | Rio Grande | West | Butte | 11.57 | https://arctos.database.museum/guid/MSB:Bird:60548 |
| Vireo | bellii | [admixed] | MSB | 280762 | 60554 | Vireo_bellii_NK280762 | NM | Rio Grande | West | Butte | 8.78 | https://arctos.database.museum/guid/MSB:Bird:60554 |
| Vireo | bellii | [admixed] | MSB | 280764 | 60556 | Vireo_bellii_NK280764 | NM | Rio Grande | East | Butte | 9.21 | https://arctos.database.museum/guid/MSB:Bird:60556 |
| Vireo | bellii | [admixed] | MSB | 280768 | 60560 | Vireo_bellii_NK280768 | NM | Rio Grande | West | Butte | 7.81 | https://arctos.database.museum/guid/MSB:Bird:60560 |
| Vireo | bellii | [admixed] | MSB | 280791 | 60583 | Vireo_bellii_NK280791 | NM | Rio Grande | West | Butte | 3.05 | https://arctos.database.museum/guid/MSB:Bird:60583 |
| Vireo | bellii | [admixed] | MSB | 279612 | 60499 | Vireo_bellii_NK279612 | NM | Rio Grande | West | Sev | 12.17 | https://arctos.database.museum/guid/MSB:Bird:60499 |
| Vireo | bellii | [admixed] | MSB | 279617 | 60500 | Vireo_bellii_NK279617 | NM | Rio Grande | East | Sev | 4.92 | https://arctos.database.museum/guid/MSB:Bird:60500 |
| Vireo | bellii | [admixed] | MSB | 279609 | 60502 | Vireo_bellii_NK279609 | NM | Rio Grande | East | Sev | 28.79 | https://arctos.database.museum/guid/MSB:Bird:60502 |
| Vireo | bellii | medius | MSB | 279618 | 60496 | Vireo_bellii_NK279618 | NM | Pecos River | East | Pecos | 33.44 | https://arctos.database.museum/guid/MSB:Bird:60496 |
| Vireo | bellii | medius | MSB | 279629 | 60497 | Vireo_bellii_NK279629 | NM | Pecos River | East | Pecos | 15.82 | https://arctos.database.museum/guid/MSB:Bird:60497 |
| Vireo | bellii | medius | MSB | 279598 | 60498 | Vireo_bellii_NK279598 | NM | Pecos River | East | Pecos | 5.52 | https://arctos.database.museum/guid/MSB:Bird:60498 |
| Vireo | bellii | medius | LSU | n/a | 47300 | Vireo_bellii_LSUB47300 | TX | W Texas | East | WTX | 12.52 | n/a |
| Vireo | bellii | medius | LSU | n/a | 47301 | Vireo_bellii_LSUB47301 | TX | W Texas | East | WTX | 9.30 | n/a |
| Vireo | bellii | medius | LSU | n/a | 51626 | Vireo_bellii_LSUB51626 | TX | W Texas | East | WTX | 8.47 | n/a |
| Vireo | bellii | medius | SDSU | n/a | 2912 | Vireo_bellii_SDSU2912 | TX | W Texas | East | WTX | 6.35 | n/a |
| Vireo | bellii | medius | SDSU | n/a | 2913 | Vireo_bellii_SDSU2913 | TX | W Texas | East | WTX | 6.42 | n/a |
| Vireo | bellii | medius | SDSU | n/a | 2914 | Vireo_bellii_SDSU2914 | TX | W Texas | East | WTX | 3.45 | n/a |
| Vireo | bellii | medius | SDSU | n/a | 2915 | Vireo_bellii_SDSU2915 | TX | W Texas | East | WTX | 7.44 | n/a |
| Vireo | bellii | medius | SDSU | n/a | 2916 | Vireo_bellii_SDSU2916 | TX | W Texas | East | WTX | 11.65 | n/a |
| Vireo | bellii | medius | UTEP | n/a | 3117 | Vireo_bellii_UTEP3117 | TX | W Texas | West | WTX | 8.06 | https://arctos.database.museum/guid/UTEP:Bird:3117 |
| Vireo | bellii | medius | UTEP | n/a | 3130 | Vireo_bellii_UTEP3130 | TX | W Texas | East | WTX | 11.82 | https://arctos.database.museum/guid/UTEP:Bird:3130 |
| Vireo | bellii | medius | UTEP | n/a | 3350 | Vireo_bellii_UTEP3350 | TX | W Texas | East | WTX | 28.76 | https://arctos.database.museum/guid/UTEP:Bird:3350 |
| Vireo | bellii | medius | LSU | n/a | 30896 | Vireo_bellii_LSUB30896 | TX | Lower RGV | East | STX | 8.00 | n/a |
| Vireo | bellii | medius | LSU | n/a | 43284 | Vireo_bellii_LSUB43284 | TX | Lower RGV | East | STX | 11.51 | n/a |
| Vireo | bellii | bellii | KU | 2707 | 89925 | Vireo_bellii_KU2707 | MO | Missouri | East | MO | 22.53 | n/a |

**Table S2:** Pairwise F_ST_ of all populations. Below the diagonal in plain text is autosome only, above the diagonal is Z chromosome F_ST_. Bolded Z chromosome values are greater than autosomal values for a given comparison and italicized values are less

| Pops | Arizona | Gila | Mimbres | Butte | Sevilleta | Pecos | W Texas | S Texas |
| --- | --- | --- | --- | --- | --- | --- | --- | --- |
| Arizona | - | **0.0215** | **0.0187** | *0.0109* | **0.0786** | **0.0853** | **0.0773** | **0.1617** |
| Gila | 0.0154 | - | *-0.0044* | *0.0029* | *0.0045* | **0.0460** | **0.0431** | **0.1202** |
| Mimbres | 0.0110 | -0.0029 | - | *0.0084* | **0.0549** | **0.0731** | **0.0703** | **0.1493** |
| Butte | 0.0130 | 0.0116 | 0.0126 | - | **0.0349** | **0.0522** | **0.0515** | **0.1213** |
| Sevilleta | 0.0314 | 0.0297 | 0.0243 | 0.0124 | - | *0.0037* | *0.0054* | *0.0301* |
| Pecos | 0.0452 | 0.0429 | 0.0422 | 0.0289 | 0.0225 | - | *-0.0040* | 0.0076 |
| W Texas | 0.0377 | 0.0363 | 0.0382 | 0.0266 | 0.0177 | -0.0015 | - | **0.0181** |
| S Texas | 0.0750 | 0.0698 | 0.0665 | 0.0506 | 0.0406 | 0.0076 | 0.0023 | - |

**Table S3:** Estimates of mean nucleotide diversity for the focal four populations, partitioned by autosome vs Z chromosome.

|  | **Arizona** | **Elephant Butte** | **Sevilleta** | **West Texas** |
| --- | --- | --- | --- | --- |
| **Autosome** | 0.0046 | 0.0045 | 0.0041 | 0.0050 |
| **Z Chrom.** | 0.0032 | 0.0032 | 0.0031 | 0.0038 |
